## Supplementary material for "Systematic exploration of domain assortments in NOD-like receptors uncovers two types of NACHT domains in *Sordariales* fungi": Figure S3

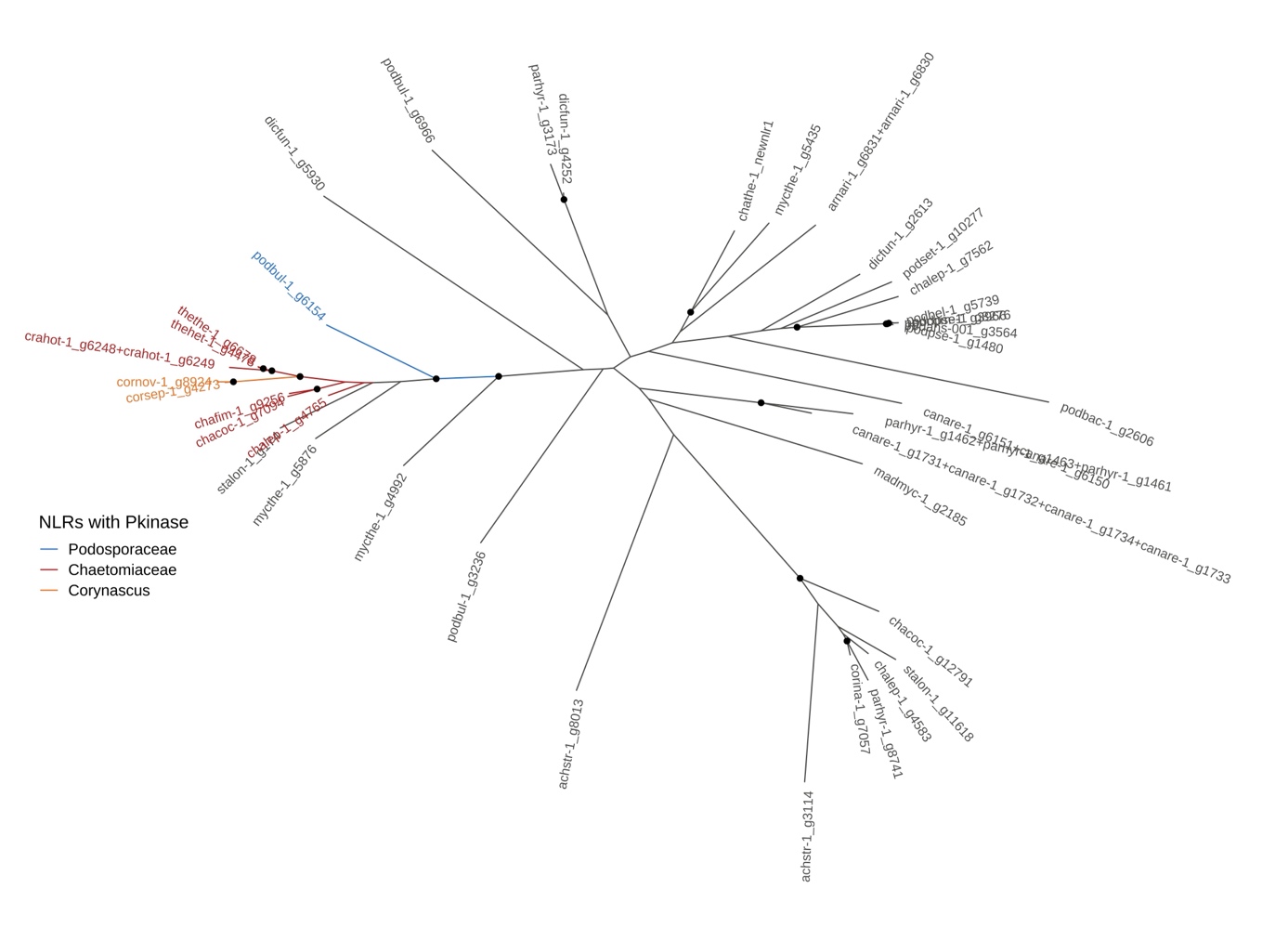

Figure S3. Phylogeny of all NACHT domains in orthogroups OGpodoTNACHT005 and OGchaetoTNACHT008. Branches corresponding to NLR sequences containing a Pkinase domain are highlighted in blue (Podospora bulbillosa), orange (Corynascus sp.), or red (other Chaetomiaceae species). Phylogeny was inferred using RAXML-NG on NACHT sequences aligned using MUSCLE. Nodes with a bootstrap value higher than 60% are highlighted with a black dot.
