## Supplementary material for "Systematic exploration of domain assortments in NOD-like receptors uncovers two types of NACHT domains in *Sordariales* fungi": Figure S2

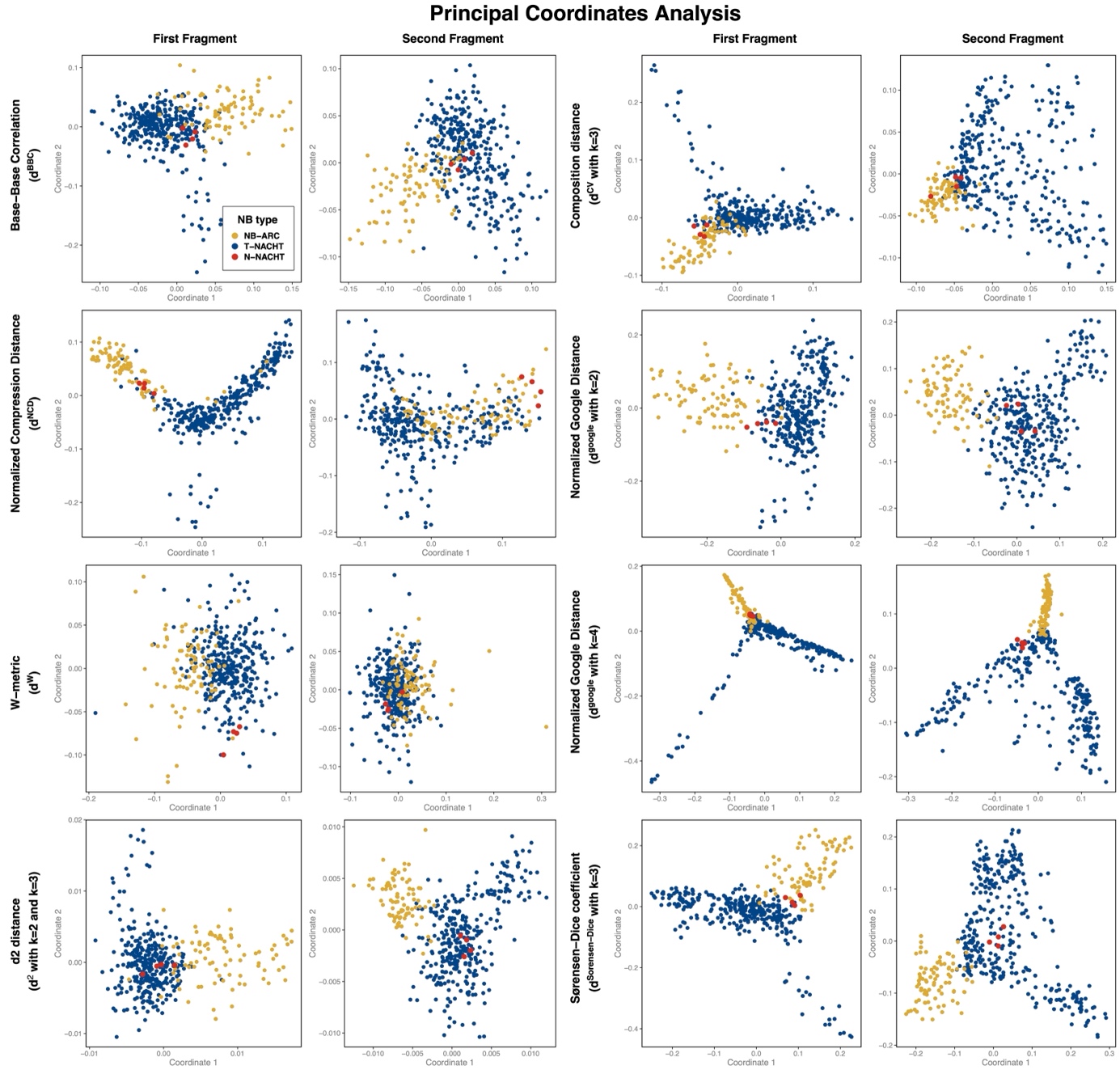

Figure S2. Principal Coordinates Analysis of distance matrices at the first and second fragments of the NB domain. The analysis was conducted with eight different modalities. Distances were computed using eight different alignment-free methods: Base-Base Correlation index (maximum distance to observe a correlation between bases set to 10), Normalized Compression Distance, W-metric (with a BLOSUM62 matrix), d2 distance (with the minimum and maximum word sizes set respectively to 2 and 3 and the vector set to frequencies), Composition Distance (with word size set to 3), Normalized Google Distance (with word size set to 2 and vector set to frequencies), Normalized Google Distance (with word size set to 4 and vector set to frequencies), Sorensen-Dice Coefficient index (with word size set to 3).
