## Supplementary material for "Systematic exploration of domain assortments in NOD-like receptors uncovers two types of NACHT domains in *Sordariales* fungi": Figure S1

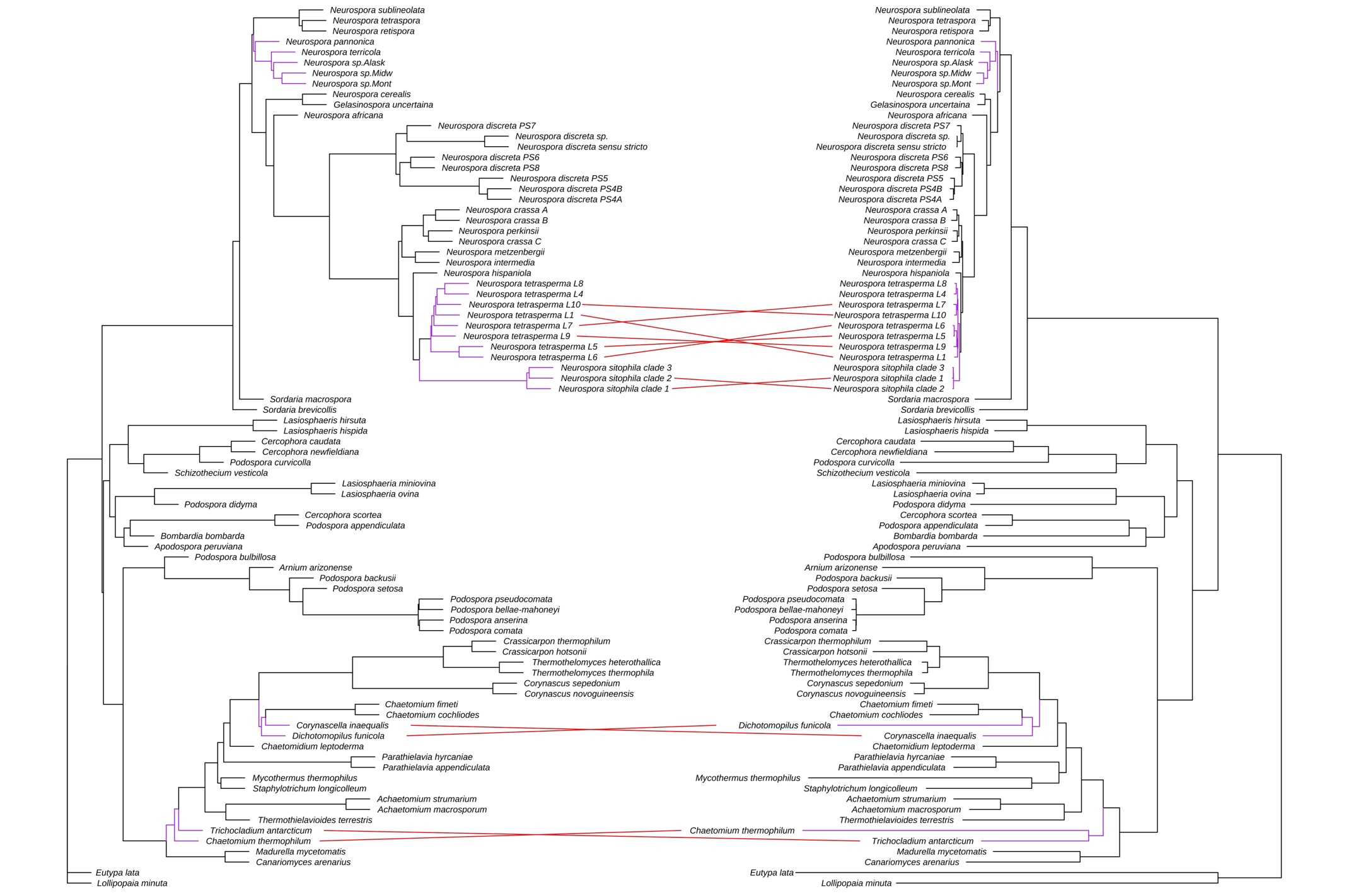

Figure S1. Comparison of species trees built using the Astral program (left) and Maximum-likelihood phylogenetic inference based on concatenated data (right). Topological differences are highlighted in purple, with the corresponding leaves connected by red lines. Astral phylogeny was built using 2367 gene trees from single-copy orthologs including at least 80% of species. Maximum-likelihood phylogeny was built using 1030 single-copy core orthologous sequences.
